# Genome characterization of ‘*Candidatus* Phytoplasma meliae’ (isolate ChTYXIII)

**DOI:** 10.1101/2021.05.31.446484

**Authors:** Franco Daniel Fernández, Luis Rogelio Conci

## Abstract

‘*Candidatus* Phytoplasma meliae’ (subgroups 16SrXIII-G and XIII-C) has been reported in association to chinaberry yellowing disease in Argentina, Bolivia and Paraguay. In Argentina, this disease constitutes a major phytosanitary problem for chinaberry forestry production. To date, no genome information of this phytoplasma and others from 16SrXIII-group has been published, hindered its characterization at genomic level. Here we analyze the draft genome of ‘*Candidatus* Phytoplasma meliae’ strain ChTYXIII obtained from a chinaberry-infected plant using a metagenomics approach. The draft assembly consists of twenty-one contigs with a total length of 751.949 bp. The annotation contains 669 CDSs, 34tRNA and one set of rRNA operons. Metabolic pathways analysis indicated that the ChTYXIII contains the complete core genes for glycolysis and functional sec system for translocation of proteins. The phylogenetic relationships inferred 132 single copy genes (orthologues core) analysis revealed that ‘*Ca*. P. meliae’ constitutes a clade closely related to the ‘*Ca*. australiense’ and ‘*Ca*. P. solani’. Thirty-one putative effectors were identified, among which a homologue to SAP11 was found and others that have only been described in this pathogen. This work provides relevant genomic information for ‘*Ca*. P. meliae’ and constitutes the first genome described for the group 16SrXIII (MPV).

## 1. Introduction

‘*Candidatus* Phytoplasma meliae’ have been associated with the yellowing disease of China tree (*Melia azedarach*) (Chinatree yellows, ChTY) in Argentina (Fernández et al., 2016) (subgroup16SrXIII-G), Paraguay (Arneodo et al., 2005) (subgroup16SrXIII-G) and Bolivia (Harrison et al., 2003) (subgroup16SrXIII-C). Chinaberry plants affected by this phytoplasma develop characteristic symptoms, such as reduced leaf size and yellowing, witches’-broom and dieback. China tree mortality caused by ‘*Ca*. P. meliae’ and Chinaberry tree decline phytoplasma (subgroup 16SrIII-B) (Galdeano et al., 2004) constitutes a serious phytosanitary problem, mainly in north-east region of Argentina, where this tree species is grown for furniture manufacturing (Arneodo et al., 2007, Fernández et al., 2015, 2016). The 16SrXIII group (*Mexican periwinkle virescence*, MPV) constitute a monophyletic clade (Figure 1) within twelve subgroups have been described so far (last one 16SrXIII-L) (Bongiorno et al., 2020). An interesting aspect of this group of phytoplasmas is their geographical distribution, which seems to be restricted to the American continent (Fernández et al., 2016; Pérez-López et al., 2016). In addition, only few host species have been associated to MPV group, among them strawberry (Jomantiene et al., 1998; Fernández et al., 2015; Pérez-López E and Dumonceaux 2016; Melo et al., 2018; Cui et al., 2019), potato (Santos-Cervantes et al., 2010), periwinkle (Lee et al., 1998), papaya (Melo et al., 2013), broccoli (Pérez-López et al., 2016) in addition to those already mentioned for ‘*Ca*. P. meliae’ above. Thanks to the reduction in the cost of sequencing in the last ten years, phytoplasma sequencing projects have increased notably. Phytoplasmas have a unique biology and genomic knowledge allowed us to understand aspects of pathogenicity and evolution never before described (Oshima et al., 2004; Chung et al., 2013; Orlovskis et al., 2017;Cho et al., 2020; Huang et al., 2021). Currently, no genomic data have been generated in any of the species that make up the MPV group, which makes it difficult to study the mechanisms associated with pathogenicity, evolution or their dispersion mediated by insects. In the present study, we report the draft genome of ‘*Ca*. Phytoplasma meliae’ strain ChTYXIII-Mo (subgroup 16SrXIII-G) obtained from infected chinaberries in Argentina. The goal of this work was to provide basic genomic information, in order to understand fundamental aspects of the evolution and pathogenicity of this phytoplasma and related phytoplasmas from MPV-clade.

**Figure 1.**
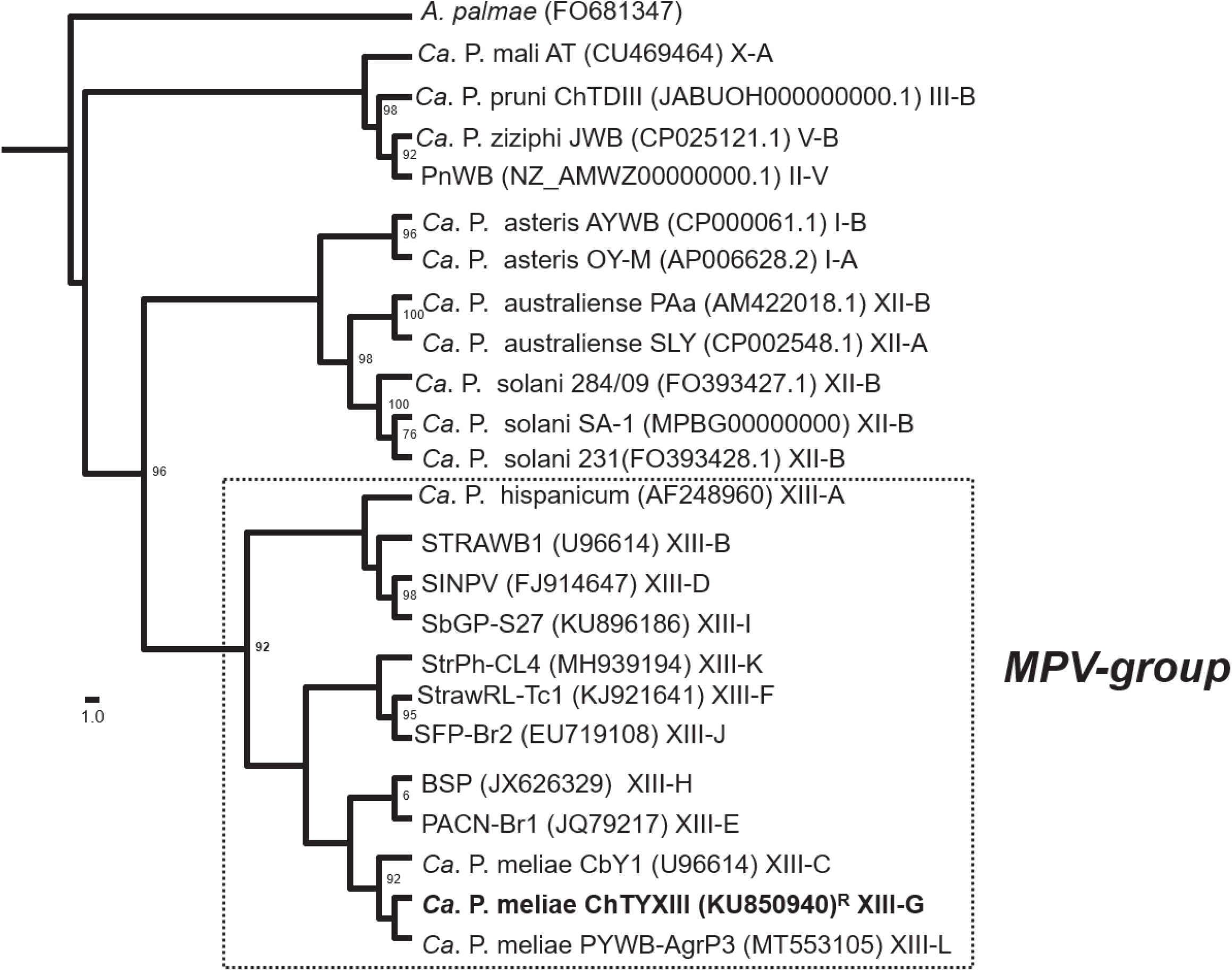
Phylogeny of phytoplasmas from group MPV based in the analysis of 16Sr RNA sequence. The tree was inferred using the maximum-likelihood method. *Acholeplasma palmae* was used as an outgroup. The numbers on the branches are bootstrap (confidence >70%) values (expressed as percentages of 1000 replicates). The GenBank accession number for each taxon is given between parentheses and the 16Sr group/subgroup classification was also provided. MPV-clade is boxed with a dotted line and ‘*Ca*. P. meliae’ strain sequenced in this study is in bold. R: reference strain.

## 2. Materials and Methods

### 2.1 Plant samples

*“Candidatus* Phytoplasma meliae” isolate ChTYXIII-Mo (KU850940) (Fernández et al., 2016) was maintained in greenhouses under controlled conditions and propagated in chinaberry tree plantlets by grafting. Total genomic DNA was extracted from infected midribs using DNeasy Plant Mini Kit (Qiagen, Germany) following the manufacturer instructions. Quality and quantity controls were assayed by electrophoresis in 1% agarose gels and spectrophotometry (Nanodrop-1000).

### 2.2 Library construction and sequencing

Total DNA was used for construction of paired-end libraries (150bp) according to TruSeq™ DNA Nano protocol and sequenced in Illumina Novaseq platform (Macrogen, Korea). Quality of RAW-reads was checked using FastQC (https://github.com/s-andrews/FastQC) and then were trimmed with Trim Galore! (https://github.com/FelixKrueger/TrimGalore) applying default settings.

### 2.3 Assembling and annotation

A metagenomic-approach was implemented for assembling based on previous pipelines with some modifications (Music et al., 2019). Trimmed reads were assembled using Unicycler (Wick et al., 2017) (Bridging mode:normal, Spades correction and Pilon polish). Assembled contigs belonging to phytoplasmas were identified by BLASTx (E=1e-20, word size=11) against a local database constructed using the 34 phytoplasma genome sequences available from NCBI (txid33926). Trimmed reads were mapped using Bowtie2 v2.3.4.3 (defaults parameters) (Langmead et al., 2012) to phytoplasma-assigned contigs. An iterative process was used until the assembly was completed. Completeness of draft-assembly was estimated by CheckM v1.0.18 (Park et al., 2015). The final draft genome was annotated using the NCBI Prokaryotic Genome Annotation Pipeline (Tatusova et al., 2016). KAAS-KEGG Automatic Annotation Server (https://www.genome.jp/kegg/kaas/) was used for functional characterization of protein coding regions and reconstruction of metabolic pathways.

### 2.4 Identification of putative effector proteins

Putative effector proteins were identified based in previous pipelines (Bai et al., 2006, Fernández et al., 2019, Music et al., 2019). Prediction of signal-peptide was conducted in Signal IP v5.0 server (https://services.healthtech.dtu.dk/service.php?SignalP-5.0) and Music et al. (2019) criteria was implemented in order to define positive candidates. Proteins which passed this filter (peptide-signal+) were analyzed in TMHMM - 2.0 server (https://services.healthtech.dtu.dk/service.php?TMHMM-2.0) and those protein without any transmembrane helices domain after signal-peptide were selected (putative secreted proteins-PSP). PSP were analyzed in the Conserved Domains Database search tool (www.ncbi.nlm.nih.gov/Structure/cdd/wrpsb.cgi, expect value = 0.01) and nuclear signal prediction was acceded using NLStradamus (http://www.moseslab.csb.utoronto.ca/NLStradamus/). The subcellular localization was accessed using the LOCALIZER (http://localizer.csiro.au/) and ApoplastP (http://apoplastp.csiro.au/) servers. The final set of proteins [signal peptide (+); trans-membrane domains outside SP (-)] were analyzed by reciprocal BLASTp searches (E-value≤1e-05) against aster yellows witches’-broom (AYWB) phytoplasma proteins (taxid:322098) for identification of SAPs homologs (Bai et al. 2009).

### 2.5 Orthologues clustering and phylogenetic analyses

Identifications of orthologous protein clusters were conducted using Orthofinder v2.5.2 (https://github.com/davidemms/OrthoFinder). Genomes sequences of representative ‘*Ca*. Phytoplasmas species were retrieved for Genbank (Table S1). For phylogenetic analyses, alignment of concatenated nucleotide sequences of single-copy core genes or single genes were constructed with MAFFT v7.450 using Geneious R.10 software (Biomatters Ltd., Auckland, New Zealand). Phylogenetic trees were constructed with IQ-TREE (http://www.iqtree.org/) (substitution model: automatic, ultrafast bootstrap=1000).

## 3. Results and discussion

### 3.1 Assembly and key features of *Ca*. Phytoplasma meliae draft genome

The genome sequencing of ‘*Ca*. Phytoplasma meliae’ (strain ChTYXIII) generated ~ 3.5Gbp of RAW reads (NCBI accession: PRJNA530090) providing a ~97X-fold coverage of draft genome, representing 21 assembled contigs totaling 751.949 bp (27.31% G+C) (Table 1). Since there are no previous estimations of chromosome size for any phytoplasmas belonging to the 16SrXIII group, we used CheckM software to evaluate the assembly quality based on the presence of conserved marker genes. According to this software, the completeness of this draft was 97.34% and possible contamination of 3.29%. Since, as has been discussed in previous works (Music et al., 2019), estimations are based on a small number of marker genes and there is little representation of phytoplasma genomes in current databases, these estimations have to be considered with caution. The 21 contigs composing the draft assembly ranged between 1.832 bp to 137693 bp (N50= 53.850). In the annotation process, 669 CDSs (full-length coding sequences) were identified, with 472 annotated as proteins with assigned function and 197 as hypothetical proteins, one operon for rRNA genes and 34 tRNAs (Table 1). Functional annotation of CDSs using BlastKOALA (https://www.kegg.jp/blastkoala/), assigned 387 of 669 CDSs (~58%) to orthologues in the KEEG database. From 299 KO categories, 240 were described with only one gene while the remaining 59 presented more than one copies representing 147 genes, ~ 21% of total CDss (49 KO with 2 genes, 4 KOs with 3 genes, 3 KOs with 4 genes, 2 KOs with 5 genes each, and a single KO to which 15 different genes were assigned). The proportion of multicopy genes in the ‘*Ca*. P. melia’ ChTYXIII genome (~ 21%) was higher than the observed in other related phytoplasmas as, ‘*Ca*. P. solani’ SA-1 (18.5%) (Music et al., 2019), ‘*Ca*. P. asteris’ strain AY-WB (10.2%), (Bai et al., 2006), OY-M (14,1%) (Oshima et al., 2004) or ‘Ca. P. australiense’ PAa (12.1%) (Tran-Nguyen et al.,2008). Multicopy genes in PMU-like regions and genome size appear to be positively correlated to a broad host range in phytoplasmas (Music et al., 2019). This is in contrast to the fact that ‘*Ca*. P. meliae’ has been only associated with two hosts, chinaberry (Harrison et al., 2003; Fernández et al., 2016) and plum (Bongiorno et al., 2020). However, the total number of hosts may have been underestimated, since there is little information regarding the presence of this phytoplasma in native species, i.e. weeds, and vector insects remain unknown.

### 3.2 Metabolic pathways

Within the three-major protein families in KEEG database, 184 CDS were assigned to *Metabolism*, 236 CDS to *Genetic Information Processing* and 54 were assigned to *Signaling and Cellular Processes* (Figure 2). Transporter membrane proteins plays fundamental roles in the phytoplasmas metabolism, allowing the incorporation of metabolites and contributing to protein secretion in the host cells cytoplasm. The ATP-binding cassette (ABC) transporters form one of the largest known protein families, and are widespread in bacteria, archaea, and eukaryotes. These proteins are best known for their role in the importation of essential nutrients and the export of toxic molecules, but they can also mediate the transport of many other physiological substrates (Davidson et al, 2008). Twenty eight genes from ‘*Ca*. P. meliae ‘genome have been described as the ABC-transporters (Table S2), including the complete pathway for spermidine/putrescine transport (potA, potB, potC and potD), lysine transport (lysX1, lysX2 and lysXY) and Zinc/Manganse/Iron (II) transport (troA, troC, troD, troB). Regarding to the protein translocation system (Sec system) we identified, *secA* (CHTY_001675), secE (CHTY_0002195), *secY* (CHTY_000200), *ffh* (CHTY_001830), *ftsY* (CHTY_001825) and *yidC* (CHTY_000350), dnaJ (CHTY_000550), dnaK (CHTY_000555), grpE (CHTY_000560) and groEL (CHTY_001355) which suggests a functional sec systems in the ‘*Ca*. P. meliae’. Within the carbohydrate metabolism, the core module for glycolysis (genes *gapA, pyk, pgk, eno, tpiA* and *gpmI*) and pyruvate oxidation (genes *pdhA, pdhB, pdhD* and *aceF*) were found, which supports that ‘*Ca*. P. meliae’ could depends on glycolysis for energy generation. This pathway was described as the major energy-yielding pathway for phytoplasmas (Kube et al., 2012). In addition, the complete ORF for gene sucrose phosphorylase (*gtfA*, CHTY_000510) was identified.

**Figure 2.**
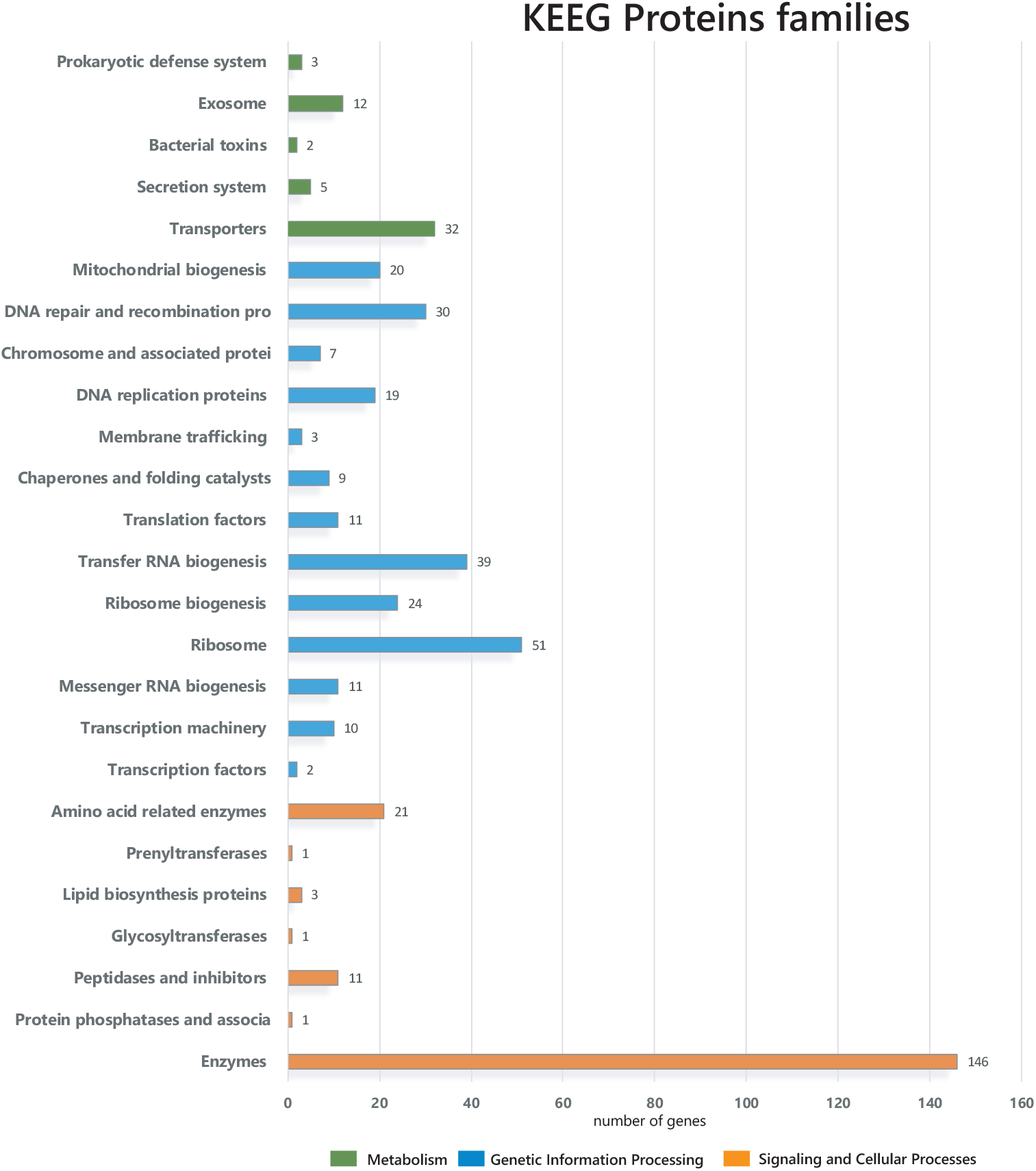
Bar chart representing the distribution of KEGG pathways associated with the draft genome of ‘*Candidatus* Phytoplasma meliae’ strain ChTYXIII. The KEGG pathways were assigned by annotating the protein coding genes using the KAAS (KEGG Automatic Annotation Server) web server.

### 3.4 Putative effectors

In the draft genome of ‘*Ca*. P. meliae’ we identified 31 proteins as putative secreted proteins (Table S3). BLASTp search against SAP-protein repertoire of AYWB phytoplasma (accession CP000061) showed the presence of putative orthologs for SAP72 (53.31%), SAP41 (52.29%), SAP21 (38.10%), SAP68 (76.00%), SAP67 (59.67%), SAP21 (38.10%) and SAP11 (50.00%). However, no homologues for SAP54, SAP05 and TENGU factor were identified. The SAP11 homologue consisted of 116 aa (CHTY_003225) with a predicted signal peptide domain (score= 0.98, position 1-32aa) and characteristic SMV signal (pfam12113, position 1-33aa/ E= 5.71e-08). Moreover, nuclear signal localization (position 35-49aa) and coiled-coiled (position 84-106aa) domains were predicted. These features are compatible with those described for the SAP11 homologues in diverse phytoplasma taxons (Bai et al., 2009; Kakisawa et al., 2014; Anabestani et al., 2017; Saccardo et al., 2012; Wang et al., 2018). A phylogenetic analysis based on amino acidic sequence for SAP11-homologues grouped the putative SAP11 protein from ‘*Ca*. P. meliae’ within the SAP11 from ‘*Ca*. P. mali’ despite they are evolutionary distant taxons (Figure 3.a). We also found a conserved synthenia in the genomic context of SAP11 with homologous regions in the chromosome of ‘*Ca*. P. asteris’ AYWB and ‘*Ca*. P. ziziphi’ JWB (Figure 3.b) suggesting possible horizontal transfer. *A. thaliana* transgenic lines that express SAP11 have curly leaves and an increased number of axillary stems that resemble the witches’ broom symptoms exhibited by AY-WB phytoplasma (Sugio et al., 2014). In greenhouse chinaberry plantlets infected by ‘*Ca*. P. melia’ shows typical witches’ broom symptoms while those infected with ‘*Ca*. P. pruni’ strain ChTDII (16SrIII-B) causes symptoms of yellowing and shortening of internodes but not witch’s broom (Figure S1). These could suggest the presence of SAP11 homologue associated mechanism in the generation of witches’ broom symptoms since no SAP11 homologues were described for ChTDIII phytoplasma (Fernández et al, 2020). A BLASTp search showed that 10 of 31 putative secreted proteins seem to be unique to ‘*Ca*. P. meliae’ since no homologs were found in the gene databank. For example, the putative protein (CHTY_003505, 226aa) present a domain associated to the TIGR04141 family, which is commonly associated to mobilomes, Others secreted protein with interesting characteristics are the CHTY_001115 (115aa), which presented a domain related to multidrug resistance efflux pump (cl34307) or CHTY_002595 (227aa) with a peptidase related domain. The protein CHTY_002535 (391aa) has a chloroplast predicted localization (score=0.991, position 105-125) and present a substrate-binding domain of an ABC-type nickel/oligopeptide-like import system (cl01709). These proteins constitute an interesting target of study as they can provide clues regarding the unique pathogenicity mechanisms of this pathogen.

**Figure 3.**
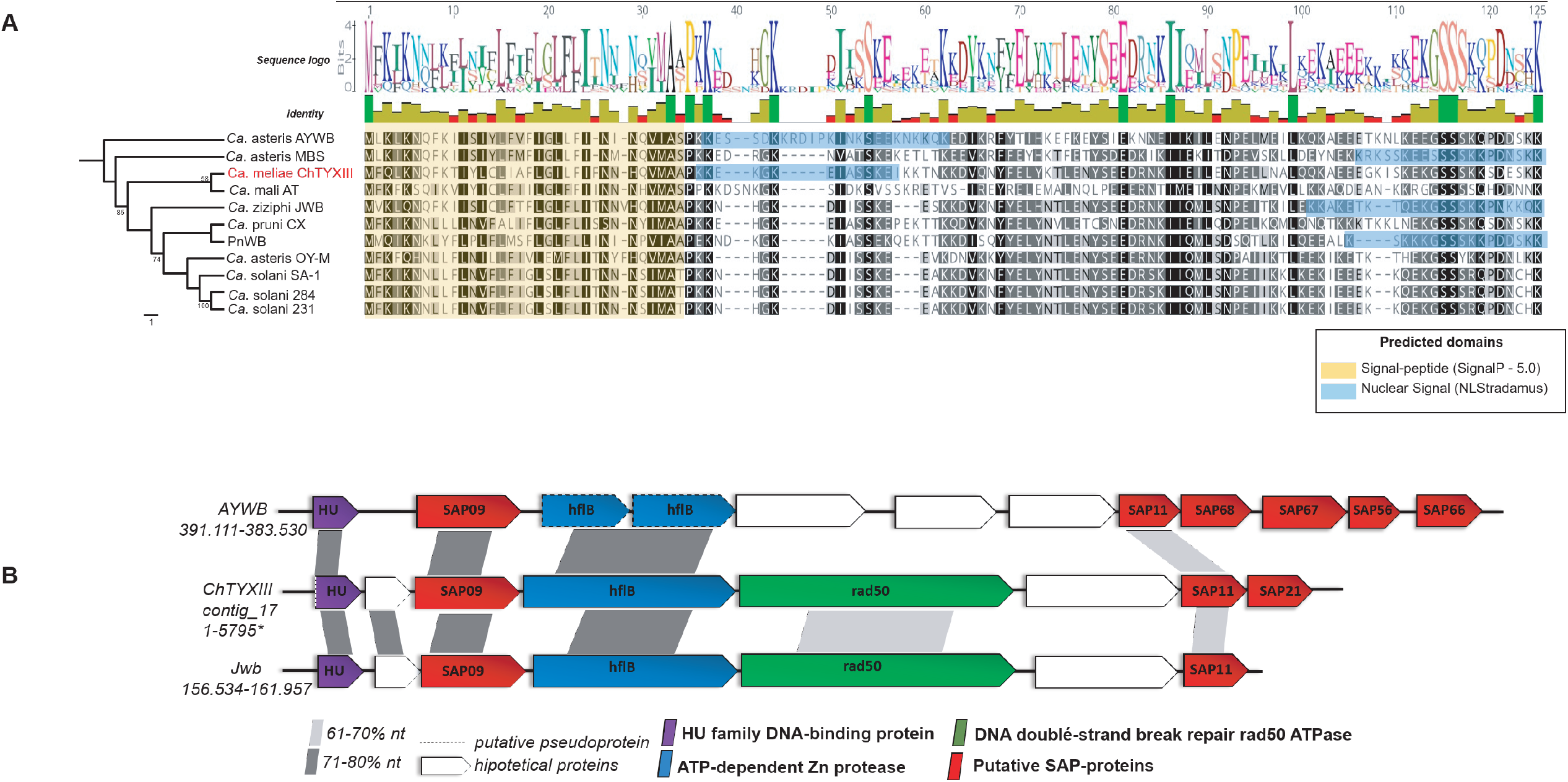
Analysis of ‘*Ca*. P. meliae’ SAP11 homologue. A: phylogeny tree inferred from aa sequence of SAP11 homologues using Maximum-Likelihood algorithm (bootstrap 1000) (scale bar: number of substitutions per site. Predicted domains (Nuclear Signal, Signal Peptide) are highlighted with color boxes in the alignment. B: synthenic organization of contig ChTYXIII-17 containing the SAP11 homologue. The genomic localization (start-end) is given below de ‘*Ca*. P. specie’ identification; SAP homologues are in red. Nucleotide sequence similarities between conserved regions are illustrated by different shades of gray colors

### 3.5 Phylogenetic analyses

Comparative genomics based on orthologues identification between draft genomes of ‘*Ca*. P. meliae’ ChTYXIII and representative genome sequence of 11 ‘*Ca*. Phytoplasma species’ (Table S1) and *A. palmae* (accession FO681347) reveals the presence of 132 single-copy genes common to all species. The phylogentic tree based on DNA and aa concatenated sequence of 132 single-copy genes showed that ChTYXIII phytoplasma form a particular clade which is more closely related to the phytoplasmas of the group 16SrXII, ‘*Ca*. P. solani strain SA-1, 284, 231 and *Ca*. P. australiense’ strain PAa and SLY (Figure 4). Similar topology was obtained when *secY* and *tufB* genes (single copy orthologs) were used (Figure S2). These results could suggest that the ChTYXIII phytoplasmas, and the associated species within the 16SrXIII group (Figure 1), could have suffered an evolutionary divergence driven by their distribution restricted to the Americas.

**Figure 4.**
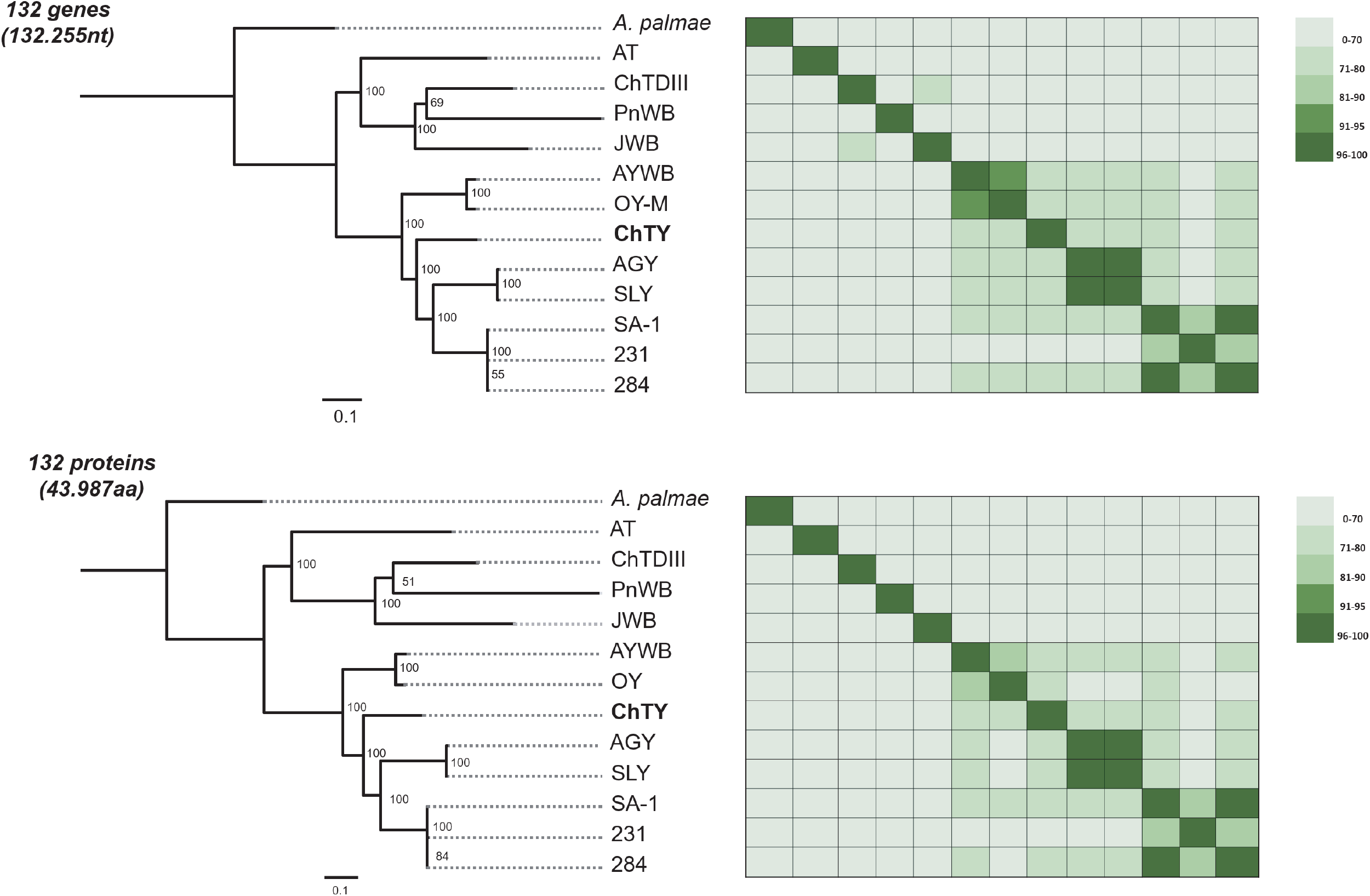
Molecular phylogeny based on nucleotide sequences and amino acid of the core genes. *Acholeplasma palmae* was included as an outgroup to root the tree. The numbers on branches indicate the level of bootstrap support (1000 replicates). The scale bar represents the number of substitutions per site. The heatmaps on the right-hand side are colored based on sequence identity.

## 4. Conclusions

This study describes the draft genome of ‘*Ca*. Phytoplasma meliae’ strain ChTYXIII. Functional analyses reveal the presence of genes related to processing and metabolism highly conserved among phytoplasmas. We have described numerous genes present in multicopy despite the fact that it is a phytoplasma with few cited hosts. Effector proteins have also been identified that could be playing key roles in the regulation of pathogenicity mechanisms. Besides SAP11 homologue, other putative effector proteins with interesting characteristics were identified, at least from the bioinformatics analysis of their domains. The phylogeny inferred from a core of genes has shown that ‘*Ca*. P. meliae’ constitutes a clade closely related to ‘*Ca*. P. solani’ and ‘*Ca*. P. australiense’. Genomic data obtained here provided new insight into the pathogenic mechanisms and evolutionary history in phytoplasmas from MPV-group.

## Funding

This research was supported by INTA, FONCyT PICT2016-0862 and PICT2017-3068. The funders had no role in study design, data collection and interpretation, or the decision to submit the work for publication.

## Data availability

RAW reads were deposited in NCBI Sequence Read Archive (SRA) under the accessions [PRJNA638346](https://www.ncbi.nlm.nih.gov/sra/14594370). The de *novo genome* draft assembly of ChTYXIII was deposited in GenBank under the accession JACAOD000000000.2 (BioProject: PRJNA63834, BioSample: SAMN15186628).

## Supplementary Material

**Figure S1.**
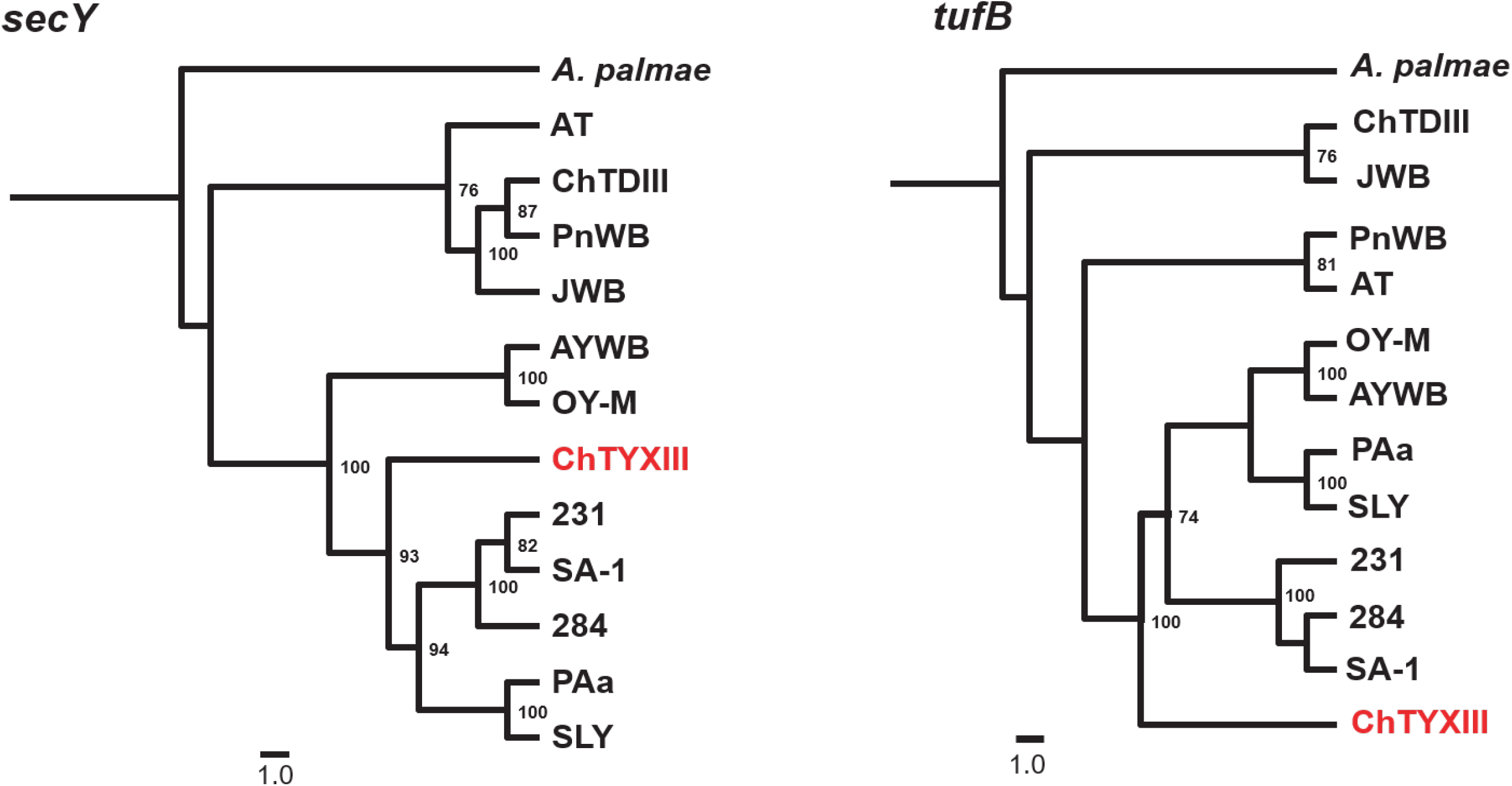
Chinaberry plantlets infected with ‘*Ca*. P. meliae’ (ChTYXIII, 16SrXIII-G) and ‘*Ca*. P. pruni’ (ChTDIII, 16SrIII-B) showing symptoms of witches’ broom and leaf yellowing and internode shortening respectively.

**Figure S2** Molecular phylogeny based on nucleotide sequences of *secY* and *tufB* genes. *Acholeplasma palmae* was included as an outgroup to root the tree. The numbers on branches indicate the level of bootstrap support (1000 replicates). The scale bar represents the number of substitutions per site.

**Table S2** List of CDSs associated to carbohydrate metabolism (Glycolysis, Pyruvate oxidation, pyruvate, Starch and sucrose metabolism), ABC transporters and bacterial sec-secretion system

**Table S3** List of putative effector proteins in the ‘*Ca*. P. meliae’ draft genome. a: presence of signal peptide (SignalP 5.0); b: localization of proteins to the plant apoplast (ApoplastP); c: subcellular localization prediction in plant cells (LOCALIZER); d: nuclear localization prediction signal (NLstradamus); e: conserved domain prediction (CDD, NCBI); f: annotation based on BLASTp analysis against ‘*Ca*. Phytoplasma’ NCBI database; g: putative SAP homologue assignment based on protein homology against SAP repertoire of AYWB phytoplasma (CP000061.1) (Bai et al., 2009). * putative proteins with no BLASTp-hit

## Notes

### Competing Interest Statement

The authors have declared no competing interest.

