## Supplementary material for "Genome characterization of ‘*Candidatus* Phytoplasma meliae’ (isolate ChTYXIII)": Table 1

|  | ***Ca*. meliae** | ***Ca*. Solani** | | | ***Ca*. australiense** | | ***Ca*. asteris** | |
| --- | --- | --- | --- | --- | --- | --- | --- | --- |
| **Features** | **ChTYXIII** | **SA-1** | **284/09** | **231/09** | **PAa** | **NZSb11** | **AYWB** | **OY-M** |
| # Contigs | 21 | 19 | 128 | 298 | 1 | 1 | 1 | 1 |
| Total Lenght (bp) | 751.949 | 821.322 | 557.538 | 515.758 | 879.324 | 959.779 | 706.569 | 860.631 |
| G+C content (%) | 27.31 | 28.30 | 28.20 | 28.60 | 27.00 | 27.20 | 27.00 | 28.00 |
| N(50) | 53.850 | 76.256 | 9.757 | 4.036 | - | - | - | - |
| assemblie status | draft | draft | draft | draft | complete | complete | complete | complete |
| total CDSs | 669 | 709 | 552 | 1146^a^ | 839 | 917 | 671 | 754 |
| CDSs w-function | 472 | 452 | 366 | 404 | 502 | 423 | 450 | 446 |
| CDSs h-protein | 197 | 257 | 157 | 169 | 337 | 494 | 221 | 308 |
| rRNA-operons | 1 | 2 | nd | nd | 2 | 2 | 2 | 2 |
| tRNA | 34 | 32 | 27 | 8 | 35 | 35 | 32 | 31 |
| Genbank # | **JABUOH010000000.2** | **MPBG01000000.1** | **FO393427.1** | **FO393428.1** | **AM422018.1** | **CP002548** | **CP000061.1** | **AP006628.2** |

**Table 1:** Genome statistic of *Ca*. Phytoplama meliae and related *Ca*. Phytoplasma species
