## Supplemental Table 1 for "Genome characterization of ‘*Candidatus* Phytoplasma meliae’ (isolate ChTYXIII)"

| ***Ca*. Phytoplasma** | **strain** | **16Sr group/subgroup** | **Host** | **Country** | **accession** | **reference** |
| --- | --- | --- | --- | --- | --- | --- |
| *Ca*. P. asteris | AYWB | 16SrI-B | *Lactuca sativa* | USA | CP000061.1 | Bai et al., 2006 |
| *Ca.* P. asteris | OY-M | 16SrI-A | *Chrysanthemum coronarium* | Japan | AP006628.2 | Oshima et al., 2004 |
| *Ca*. P. australiense | PAa | 16SrXII-B | *Catharanthus roseus* | Australia | AM422018.1 | Tran-Nguyen et al., 2008 |
| *Ca*. P. australiense | [PYL](https://www.ncbi.nlm.nih.gov/genome/1752?genome_assembly_id=171613) | 16SrXII-B | *Fragaria* x *anannassa* | New Zealand | CP002548.1 | Andersen et a., 2013 |
| *Ca.* P. mali | AT | 16SrX-A | *Catharanthus roseus* | Germany | CU469464.1 | Kube et al., 2008 |
| Ca. P. ziziphi | Jwb-nky | 16SrV-B | *Ziziphus jujuba Mill.* | China | CP025121.1 | Wang et al., 2018 |
| *Ca.* P.s pruni | ChTDIII | 16SrIII-B | *Melia azedarach* | Argentina | JABUOH000000000.1 | Fernandez et al., 2020 |
| *Ca*. P. solani | SA-1 | 16SrXII-A | *Vitis vinifera* | Italy | MPBG00000000 | Music et al., 2019 |
| *Ca*. P. solani | 231 | 16SrXII-A | *Petroselinum sativum* | Serbia | FO393428.1 | Mitrovic et al., 2014 |
| *Ca*. P. solani | 284 | 16SrXII-A | *Nicotiana tabacum* | Serbia | FO393427.1 | Mitrovic et al., 2014 |
| no assigned | PnWB | 16SrII-V | *Arachis hypogaea* | Taiwan | NZ_AMWZ00000000.1 | Chung et al., 2013 |
| ***Ca*. P. meliae** | **ChTYXIII** | **1SrXIII-G** | ***Melia azedarach*** | **Argentina** | **NZ_JACAOD000000000.2** | **This paper** |

**Table S1:** List of reference genomes used in this study. In bold draft genome obtained in this paper
